# Leaf—An open-source, model-agnostic, data-driven web application for cohort discovery and translational biomedical research

**DOI:** 10.1101/632471

**Authors:** Nicholas J Dobbins, Clifford Spital, Robert Black, Jason Morrison, Bas de Veer, Liz Zampino, Robert Harrington, Bethene Britt, Kari Stephens, Adam Wilcox, Peter Tarczy-Hornoch, Sean D. Mooney

## Abstract

**Objective:** Academic medical centers and health systems are increasingly challenged with supporting appropriate secondary use of data that originate from multiple sources. Enterprise Data Warehouses (EDWs) have emerged as central resources for these data, but they often require an informatician to extract meaningful information, thereby limiting direct access by end users. To overcome this challenge, we have developed Leaf, a lightweight self-service web application for querying and extracting clinical data from heterogeneous data sources.

**Materials and Methods:** Leaf utilizes a flexible biomedical concept system to define hierarchical items and ontologies. Each Leaf concept contains both textual representations and associated SQL query building blocks, exposed by a simple drag-and-drop user interface. Leaf generates abstract syntax trees which are compiled into dynamic SQL queries.

**Results:** Leaf is a successful production-supported tool at the University of Washington, which hosts a central Leaf instance querying an EDW with over 300 active users. Through the support of UW Medicine (https://uwmedicine.org), the Institute of Translational Health Sciences (https://www.iths.org) and the National Center for Data to Health (https://ctsa.ncats.nih.gov/cd2h/), Leaf source code has been released into the public domain at https://github.com/uwrit/leaf.

**Discussion:** Leaf allows the querying of single or multiple clinical databases simultaneously, even those of different data models. This enables fast installation without costly extraction or duplication from existing databases.

**Conclusion:** Leaf differs from existing cohort discovery tools because it does not specify a required data model and is designed to seamlessly integrate with existing enterprise user authentication systems and clinical databases in situ. We demonstrate its unique technical strengths and success alongside its friendly user interface. We believe Leaf to be useful for health system analytics, clinical research data warehouses, precision medicine biobanks and clinical studies involving large patient cohorts.

## Introduction

Healthcare organizations are challenged to manage ever increasing quantities of data, driven by the rise of electronic health record (EHR) systems, patient-reported outcomes, registries, *m*Health devices, genomic databases, and other clinical and non-clinical systems^1, 2, 3^. Beyond importing these heterogeneous and complex data into EDWs, healthcare organizations are responsible for cleaning, integrating, and making the data accessible to appropriate users and consumers, maintaining organizational compliance, and protecting patient privacy and preventing security breaches. Even in cases where data are successfully integrated and made available, their extraction can be time-consuming and difficult for consumers who may not have IT or informatics backgrounds.

To advance the needs of our health system at the University of Washington (UW) and potentially other Clinical and Translational Science Awards (CTSA) centers, we developed Leaf, a lightweight drag-and-drop web application that enables direct querying of arbitrary clinical databases. Because Leaf is a ‘blank canvas’ and does not require a particular clinical data model, making new data sources and biomedical concepts available in Leaf is often relatively simple, with no data extraction or transformation required. Utilizing an Agile development process with ongoing engagement of clinical users and stakeholders, we have incrementally improved Leaf functionality based on user feedback, adding capabilities such as real-time de-identification algorithms and direct REDCap^4^ export. We have prioritized the use of human-centered design^5^ techniques and modern web best practices to make Leaf’s user interface clear and intuitive. The result is a simple but powerful tool for our clinicians and researchers that requires fewer technical resources to maintain compared to other cohort discovery tools.

## Background and Significance

Cohort discovery tools have been demonstrated to be extremely useful for observational research, clinical trial recruitment, and hospital quality improvement programs ^6, 7, 8, 9^. In order to find which cohort discovery tools were best suited to our institution, we investigated several, including Informatics for Integrating Biology and the Bedside (i2b2)^6^ and TriNetX^10^. Our aim was to find a tool that was lightweight and easily deployable, adaptable to new data sources and use cases, and intuitive for our clinical and research users. As our institution operates a central EDW that is jointly used for quality improvement, analytics, and research, we also sought to avoid creating a costly data extraction process that would duplicate our EDW, instead working to leverage the EDW itself. The opportunities we perceived which drove the development of Leaf are summarized below.

### Opportunity 1: A lightweight ‘sidecar’ data service to the EDW

Data sources, schema, and row values in our EDW evolve with our EMRs and new institutional use cases require new sources be ingested. Often this evolution of data cannot be predicted far ahead of time, and any tools using EDW data must therefore be flexible and adaptable, ideally with minimal informatics effort. Our institution also made a significant investment in our EDW as a valuable resource available across our enterprise, and we sought a tool that did not attempt to recreate functionality in our EDW but instead leveraged and complemented it. Finally, we envisaged a modular, modern, cloud-friendly tool that could be effectively be deployed in either cloud architectures or on-premises, interoperating alongside other tools in a service-oriented fashion.

### Opportunity 2: Customized user interface for different sites/data

A second opportunity was the need for an intuitive user interface that could accommodate a wide array of data sources in a flexible manner and facilitate discoverability without overwhelming or frustrating users. We had learned from experience that an effective user interface needed to allow for both ontology-driven hierarchical data elements, such as diagnosis and procedure codes, and also local data presented in unique structure and textual descriptions. We further aimed to design an interface that could display not only aggregate patient counts, but also simple demographic visualizations and row-level cohort data available in a single click.

### Opportunity 3: Better alignment with health system analytics

A third opportunity, related to the structure of UW Medicine informatics teams, was to reduce duplication of efforts between our research informatics team and other informatics teams involved in hospital operations and analytics. As each of these teams uses our EDW to query, catalogue, and extract data, we perceived a unique opportunity to design a tool which served varied but often highly similar use cases for each. By allowing users to simply indicate their purpose (quality improvement or research) and de-identify data as needed, we believed each informatics team and their users could greatly benefit.

Leaf began as a small innovative project by the first author to create a user-friendly tool that matched the powerful capabilities of the i2b2 web application but queried our EDW directly, leveraged the existing work of teams across our organization, and functioned with no data extraction required. After an initial proof-of-concept application was developed, Leaf was shown to leaders within our hospital system and CTSA, eventually becoming a flagship collaborative software project to improve self-service access to clinical data across research, analytics, and biomedical informatics academic department at the University of Washington.

## Materials and Methods

### Agile development and user engagement

Early in Leaf development we sought to utilize Agile development methodology^11^ to ensure the Leaf user interface and features were driven by the needs of clinicians and researchers. The Leaf pilot project was formed to engage approximately 30 clinicians and researchers across various medical disciplines and organizations in our CTSA in weekly Agile sprints over approximately six months.

After an initial meeting was held to introduce the tool and discuss the scope of the project, weekly web-based meetings were conducted to iteratively introduce new features, gather feedback, and prioritize items for upcoming sprints. After each sprint, the development team worked to incorporate stakeholder ideas and coordinate with our EDW team to make additional data elements available.

After the pilot phase was complete, we planned and created a multi-tiered support system to triage user assistance and provide oversight of future Leaf development. In February of 2018, a central instance of Leaf querying our EDW was approved as an institutionally-supported tool at the University of Washington.

**Figure 1.**
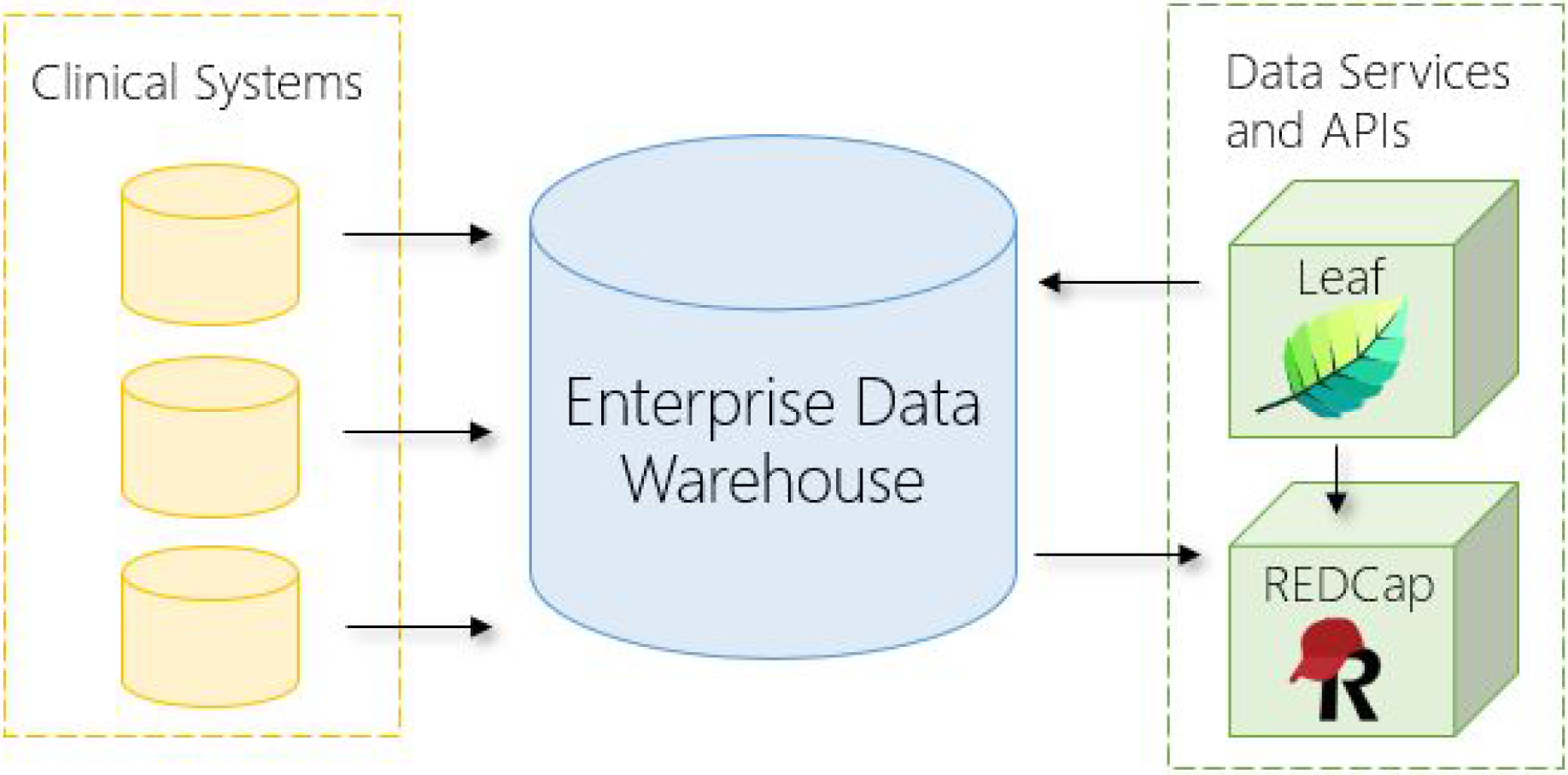
Diagram of the Leaf instance deployed at the UW Medicine Enterprise Data Warehouse. Data from EMRs and other clinical systems flow into the EDW. Leaf, REDCap, and other tools are deployed in modular fashion to query the EDW and interoperate for data extraction and analysis.

### System Architecture

A Leaf instance is typically deployed with a 1:1 relationship to the clinical database or database server which it is intended to query (hereafter for simplicity we refer to clinical database in the singular, but multiple clinical databases on a single server can be queried in the same way provided that they share a Master Patient Index^12^). A Leaf instance is composed of three core architectural elements common to many modern web applications:

- A user-facing client application written in React and TypeScript.
- A RESTful^13^ web application programming interface (API) deployed to a web server written in C# and .Net Core.
- A small SQL Server application database for logging activity, tracking queries, and caching patient identifiers.

Additionally, a separate web server is typically used to filter web traffic and serve the client application to users in coordination with a SAML2-compliant identity provider, such as Shibboleth or ADFS. All components can be deployed on either Linux or Microsoft Windows operating systems.

### Data model assumptions

Leaf makes minimal assumptions about any clinical database it queries. Indeed, one of the key insights of the Leaf application is that nearly all standard and non-standard clinical databases conform to certain simple assumptions, which makes the largely model-agnostic nature of Leaf possible. The assumptions are:

1. The clinical database to be queried has a common identifying column name for patient identifiers across tables, such as `PersonId, patient_id`, or `PAT_NUM`.
2. Similarly, the database has a common identifying column name for clinical encounter identifiers, such as `EncounterId` or `visit_occurrence_id`.

These columns are defined globally for a given Leaf instance, and are expected to be present on every table, view, or subquery to be queried, though encounter identifier columns need only be present if a given SQL set has a 1:M relationship for a patient. In tables where these columns are not present or are named differently, SQL views or subqueries can be used instead to derive them.

### Access and De-identification

Leaf is designed to be able to query clinical databases with identified patient information^14^ and hide or obfuscate that information when users access Leaf in ‘de-identified mode’. The workflow for this is:

1. On login users are presented with options for accessing data for the purposes of quality improvement or research. They must then enter quality improvement documentation (i.e., project name, approving body) or institutional review board (IRB) approval information, or alternatively indicate that they do not have this information. Users then select identified or de-identified access, with access limited to de-identified mode if the user has no approved quality improvement project or IRB.
2. As users run queries and view row-level data in the ‘Patient List’ screen, the Leaf REST API de-identifies the output SQL data depending on the use type. If Leaf is in de-identified mode, SQL date-type field values are shifted forward or backward a number of hours based on a randomly-generated value for each patient unique to the query the patient was included in, while HIPAA identifier fields are removed before sending results to the client.

For information on Patient List datasets based on FHIR templates, see the *Extracting and exporting data for a cohort* section below.

In addition to the above workflow, at our institution Leaf has IRB approval for use in querying clinical data in our EDW. We further audit user-entered IRB and usage information using Leaf log files to check for inappropriate use.

### Concepts and Query compilation

The Leaf application database has a central SQL table (`app.Concept`) which defines the concepts that comprise Leaf queries, their hierarchical relationship to other concepts, and textual representations seen by users. Each concept also contains a field for an arbitrary SQL `WHERE` clause which functions as a programmatic representation of the concept in the clinical database to be queried (example in Figure 2). The SQL `FROM` clause and corresponding date field (if any) for concepts are similarly found via a foreign key relationship to the `app.ConceptSqlSet` table, and thus all relevant SQL data for a concept can be returned in a simple `JOIN`.

**Figure 2.**
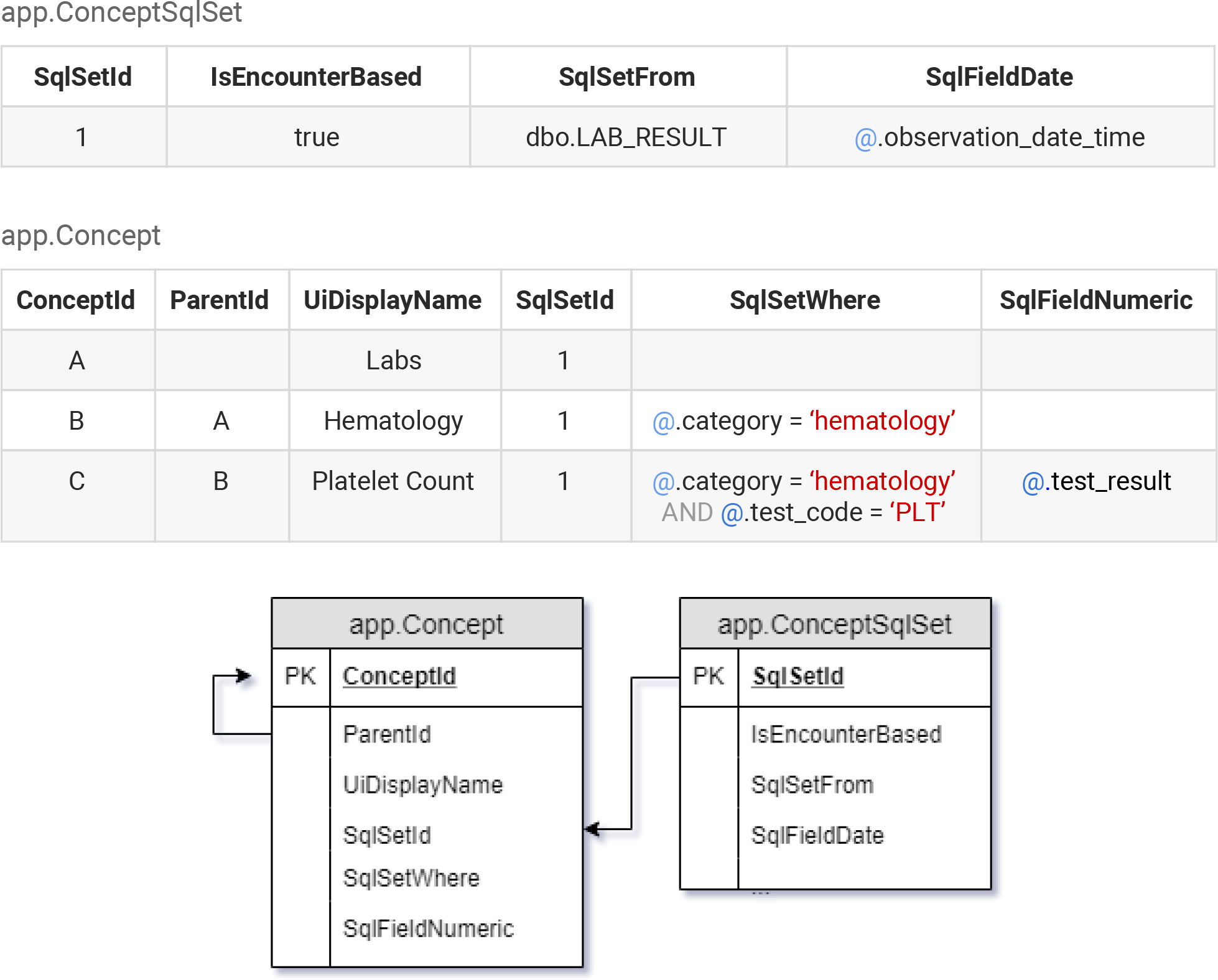
Example tabular values and Entity-Relation diagram illustrating hypothetical Leaf concepts for Laboratory tests, Hematology-based tests, and Platelet Count tests. Each concept has both textual and programmatic representation components. The ‘@’ symbols within the `SqlSetWhere` field denote ordinal positions to insert the SQL Set alias at query compilation time. Additional fields in the tables have been omitted for brevity.

This structure allows concepts to be remarkably flexible in terms of programmatic representation (Figure 2) and presented visually to users in an intuitive fashion (Figure 3). Leaf’s concept table therefore functions as a programmatic and ontological metadata map of queries to a clinical database.

**Figure 3.**
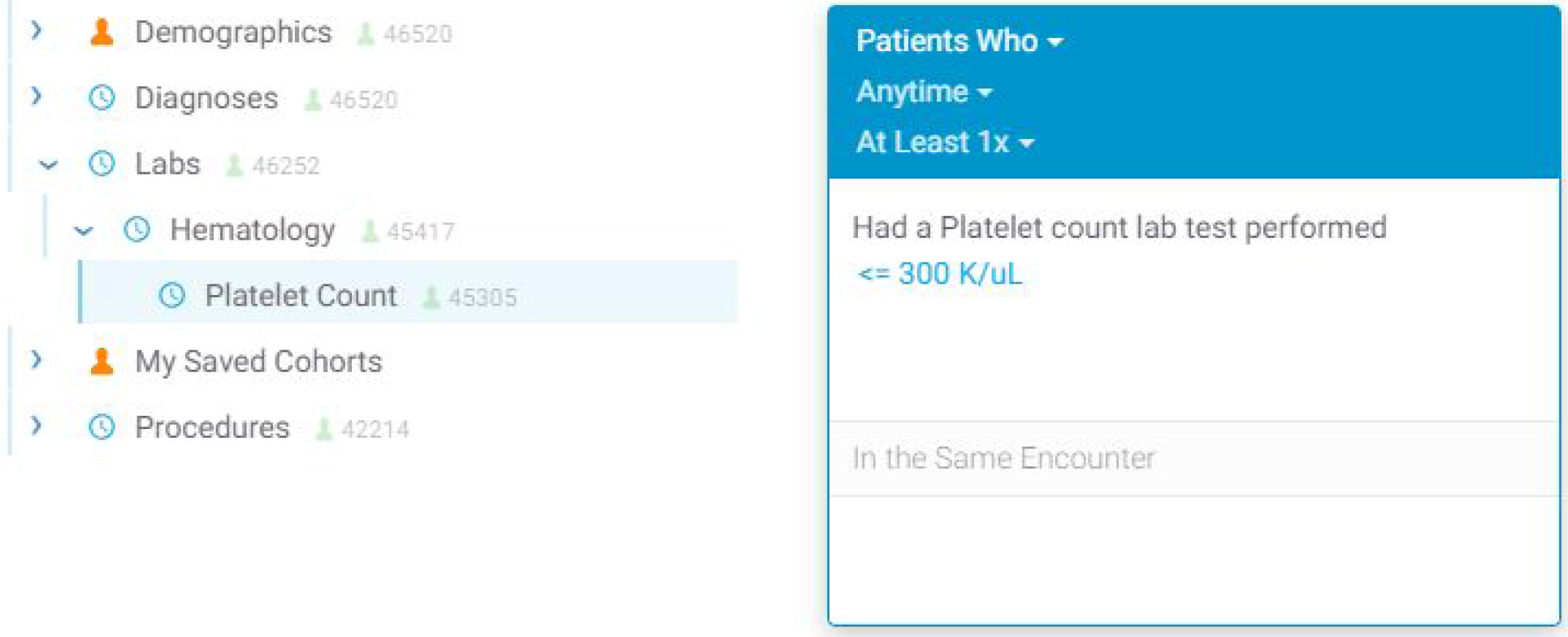
User interface screenshots of the Platelet Count example concept as seen in Figure 1. The left image shows the concept within an intuitive nested hierarchy below ‘Hematology’ and ‘Labs’, while the right image shows a simple visual query representation created after the user has dragged over the ‘Platelet Count’ concept and filtered by a value (<=300 K/uL).

To date, Leaf instances have been deployed and tested within our development environments and we have successfully validated this approach with a variety of data models, including the proprietary vendor-specified model of our EDW, the OMOP Common Data Model^15^, i2b2^6^, Epic Clarity^16^, and MIMIC-III^17^.

### Query compilation and concept mapping

As users drag concepts over to define queries, the Leaf client creates an array of abstract syntax tree (AST) objects in JavaScript Object Notation (JSON) representing the current user interface state and query definition. The AST objects themselves do not contain SQL, but instead contain `ConceptId` pointers to concepts in the Leaf `app.Concept` table included in the user’s query, along with additional metadata describing the query.

When the AST objects are sent to the Leaf REST API after the user clicks ‘Run Query’, the API performs the following steps:

1. Validates that the user is permitted to use each requested concept in the AST based on the `ConceptId` values in the Leaf application database.
2. Loads the SQL `FROM` and `WHERE` clauses and appropriate date and numeric fields (if applicable) for each concept.
3. Generates individual SQL queries for each of the 3 ‘panels’ in the Leaf user interface that can be used to create queries. Empty panels are excluded.
4. Computes a summed estimated query cost per panel to the database using a simple heuristic determined by cached patient counts per concept as proxy for cost, with a reduction if concepts are constrained by dates or numeric values.
5. Generates a SQL Common Table Expression (CTE) by ordering the panels by the estimated query cost values generated in (4). The queries are then linked by SQL `INTERSECT` and `EXCEPT` statements starting with the least estimated cost to optimize performance.
6. Runs the query against the clinical database, map-reducing the resulting patient identifiers in a hashset and caching them in the Leaf application database should the user later request row-level data for the cohort.
7. Returns a count of unique patients to the client application.

### Building the concept tree using ontologies and the UMLS

Because Leaf concept definitions are stored in a simple SQL table, relational database-based ontological mapping systems such as the Unified Medical Language System^18^ (UMLS) can be leveraged to dynamically build the Leaf concept tree programmatically, as has been demonstrated with i2b2^19^. The UMLS has been used extensively in our central EDW Leaf instance to build the concept tree sections for diagnosis and problem list codes (ICD-9, ICD-10, and SNOMED), procedure codes (HCPCS, CPT, CDT, ICD-9, and ICD-10), and laboratory tests (LOINC).

SQL-generated Leaf concepts can also be data-driven and unique to local clinical data. Scripts to generate unique concepts for clinics, services, hospital units, demographic characteristics, and other data types can be used to create arbitrary institution-specific concepts. Data such as those derived from natural language processing can similarly be made available in this way.

### Querying multiple Leaf instances simultaneously

Because Leaf AST-based query representations do not contain SQL but rather pointers to arbitrary concepts, mapping Leaf queries to various data models is notably straightforward, with a few important caveats.

Each Leaf concept contains an optional `UniversalId` field for the purpose of mapping the concept to a matching concept in a different Leaf instance, provided that the `UniversalId` values are the same. Concept `UniversalId` values must conform to the Uniform Resource Name (URN) specification^20^ and begin with the prefix `urn:leaf:concept`, but otherwise have no restrictions.

For example, two or more institutions with Leaf instances installed can choose to allow their users to query each other’s clinical databases using Leaf. After exchanging secure certificate and web address information, administrators must agree on common naming conventions for Leaf concept `UniversalIds`. A `UniversalId` of a concept for ICD-10 diagnosis codes related to type 2 diabetes mellitus could be defined as

~~~
urn:leaf:concept:diagnosis:coding=icd10+code=e11.00−e11.9
~~~

which serves as a human-readable identifier and includes the domain (diagnosis), the coding standard (ICD-10), and relevant codes to be queried (a range of diabetes mellitus type 2-related codes from E11.00 to E11.9).

Users from each institution continue to see the concepts defined by their local Leaf instance in the user interface. As users run queries across partner Leaf instances, however, each partner instance automatically translates the sender’s concepts to local concepts by `UniversalIds`. After concept translation, local instance-specific SQL queries are generated and executed, with results returned to the user. If a given Leaf instance does not have a local concept for any one of the concepts included in the user’s query, it responds with that information to the user and does not create a query.

Because each Leaf instance translates and runs queries independently, a missing concept in one instance does not affect others, which proceed to report results to the user.

### Boolean logic and temporality constraints in cohort definition queries

Leaf’s core functionality is as a flexible query constructor and execution engine. Queries can be configured to handle `AND`, `OR`, negation, and various temporality constraints, and include multiple visual cues to users. Temporal query relations are created vertically in the user interface (Figure 4), while additional inclusion or exclusion criteria with no temporal relations are represented horizontally in multiple panels (Figure 5). The query constructor user interface is designed to approximate natural language as much as possible in order to make query logic clear and readable to users.

**Figure 4.**
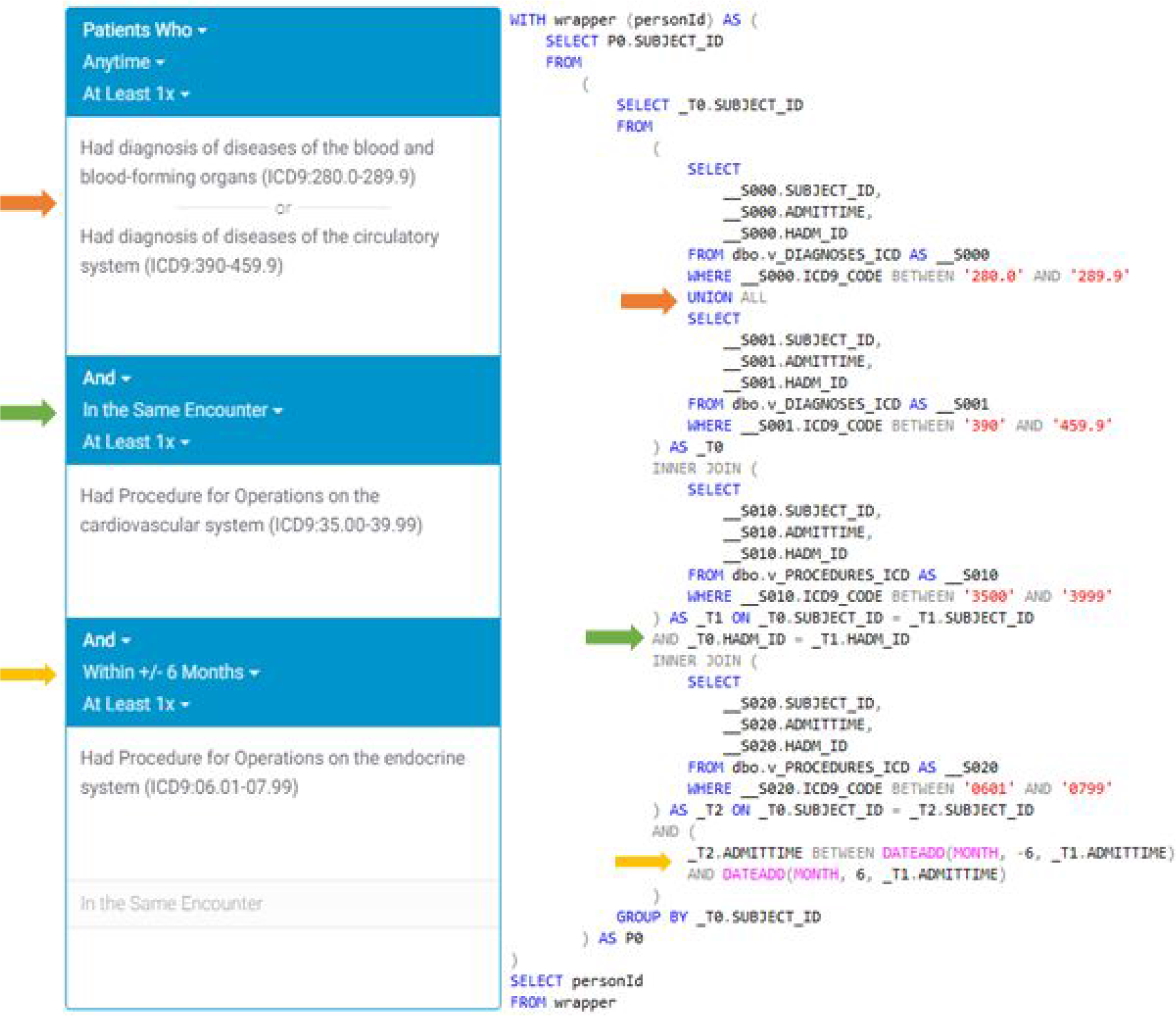
Visual representation of a single-panel query with the user interface on the left and corresponding compiled SQL representation on the right using the MIMIC-III database^17^. The example demonstrates the use of OR, Same Encounter, and temporal range logic.

**Figure 5.**
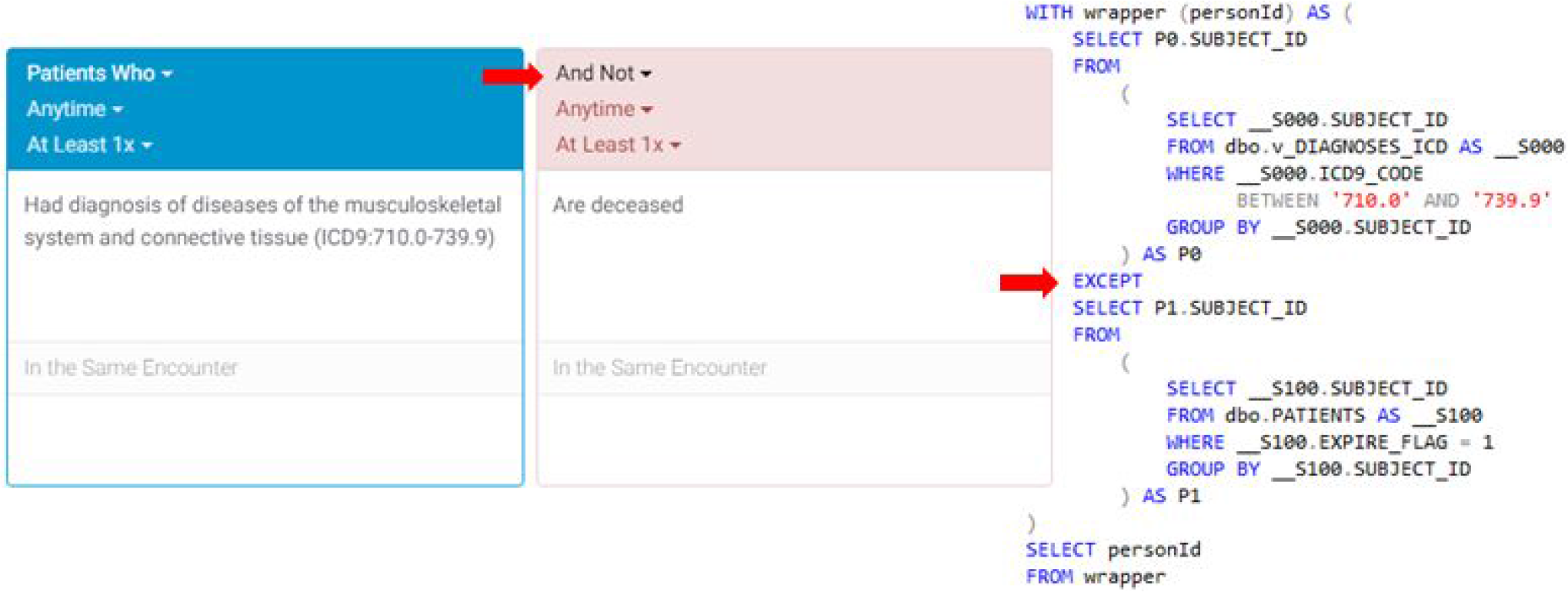
Visual representation of a two-panel query with the user interface on the left and corresponding compiled SQL representation on the right. The second panel is used as an exclusion criteria and executed using a SQL `EXCEPT` statement.

### Extracting and exporting data for a cohort

After executing a cohort estimation query, Leaf allows users to explore row-level data using the ‘Patient List’ screen. Exportable datasets (hereafter simply ‘datasets’) are defined by an administrator and configured by dynamic SQL statements whose output must conform to a template of predefined column names. Dataset templates include ‘Encounter’, ‘Condition’, ‘Observation’, ‘Procedure’, and ‘Immunization’, and are based on denormalized tabular representations of FHIR resources^21^.

Like concepts, Leaf datasets include an optional `UniversalId` field which allows multiple Leaf instances to return predictable row-level FHIR-like clinical data (Figure 6), even if their underlying data models differ. After data are returned, users can quickly create and populate custom REDCap projects through the Leaf interface in both an identified and de-identified fashion. Leaf also employs an administrator-defined row export limit to ensure smooth operations and prevent large dataset exports without proper oversight.

**Figure 6.**
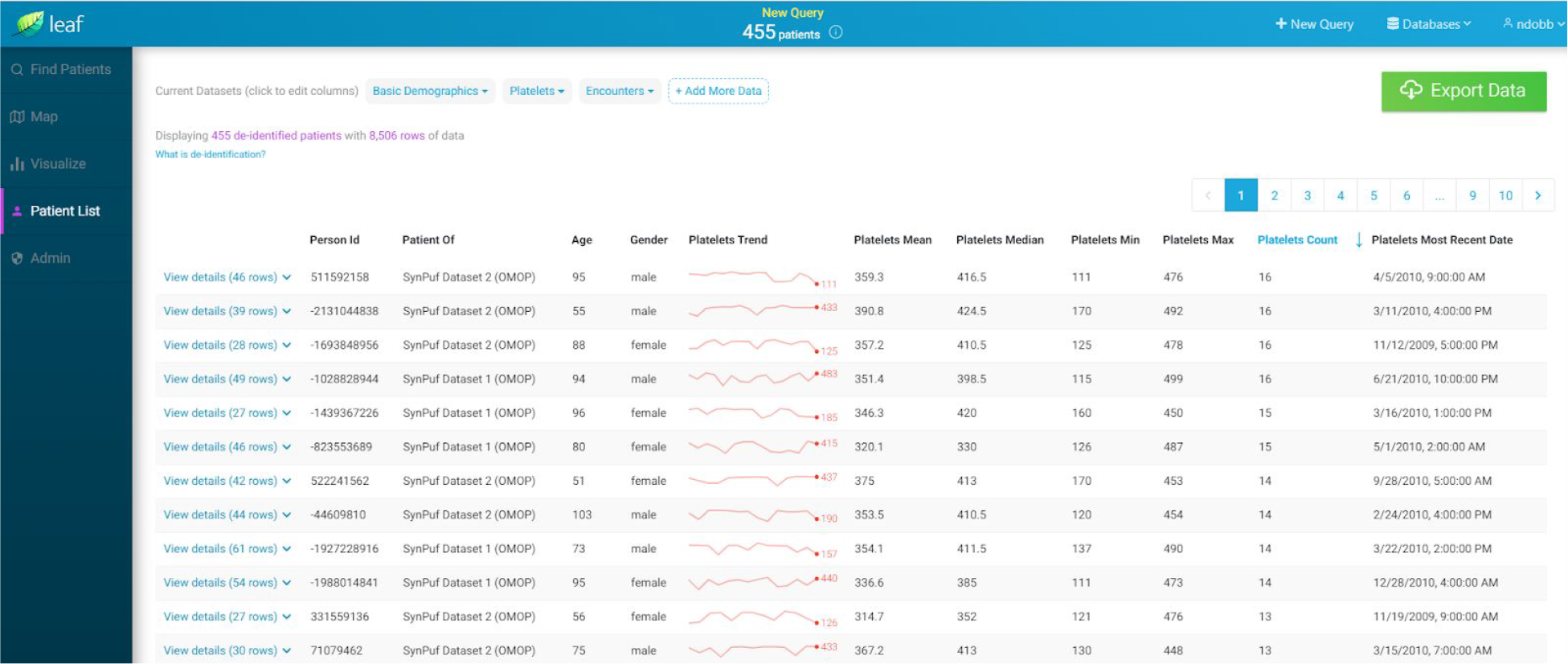
Screenshot of the Leaf user interface showing synthetic demographic and laboratory result data for a cohort from multiple Leaf instances, querying OMOP and i2b2 databases simultaneously. The Leaf client application automatically generates summary statistics for each patient. Granular row-level data can be accessed by clicking on a patient row in the table, and is directly exportable to REDCap via the REDCap API.

## Results

Since being released as a production tool at the University of Washington in 2018, Leaf users have run over 18,000 queries against our EDW over approximately 14 months (Table 1). During this time, our research informatics team has noted a relative increase in complex data extraction requests as usage of Leaf has grown, with a corresponding decrease in requests for simpler queries which we believe are likely being accomplished by clinicians and researchers using Leaf, though this is difficult to directly measure.

**Table 1.**
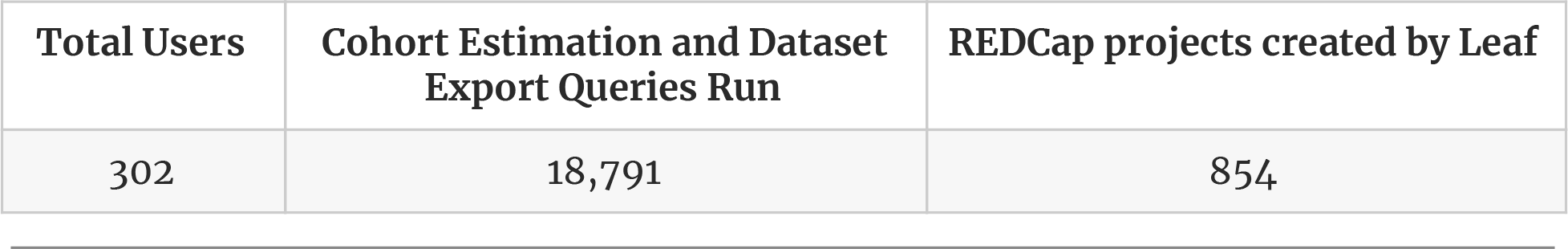

Information on Leaf is available within our institution on the Institute of Translational Health Science’s website, internal intranet sites at the University of Washington, and through introductory presentations we have conducted. Many users continue to learn of Leaf by word of mouth, however, as we’ve pursued a ‘soft’ production release in order to estimate informatics resources to support a growing user base going forward. Since going live, users have shared their findings and uses of Leaf with us, including published manuscripts and abstracts^22–32^, presentations^33–35^, a summer internship research program^36^, power analysis for grant submissions, quality improvement projects, clinical trials screening, and providing local context to national studies^37,38^.

## Discussion

Leaf represents a significant advancement in data-driven self-service cohort discovery tools by allowing CTSAs, large clinical research projects and enterprises to leverage existing clinical databases while reducing the burdens of data extraction and query construction on informatics teams. Because Leaf is a self-service tool, it removes the ‘informatics barrier’ that many clinicians, trainees and students face when trying to launch observational research or quality improvement initiatives and uncorks the potential of these faculty, fellows, residents and students to implement their ideas.

Since becoming a formal project within the Center for Data to Health, we have refined Leaf to be readily accessible and valuable to the broader CTSA community. We are currently working with a number of CTSA sites to assist in piloting Leaf at other institutions.

## Limitations

Despite its strengths, Leaf has a number of limitations that should be recognized. In terms of federated queries to multiple Leaf instances, the `UniversalId` model as a means of cross-institutional querying can require significant curation by administrators to ensure that the concept `UniversalIds` at each site match exactly, though this saves the costly effort of transforming and aligning data models. Also, while designed to be intuitive, the Leaf user interface requires some training to become proficient; many University of Washington users have requested assistance in crafting queries, even after receiving training. We suspect this is caused by the complexity of the underlying clinical data instead of the complexity of Leaf. Finally, while Leaf allows for highly flexible SQL queries, it is not designed to clean, integrate, or ensure semantic alignment among data, and is not a substitute for proper database maintenance and refinement.

## Conclusion

Leaf represents a successful example of a flexible, user-friendly cohort estimation and data extraction tool that can adapt to nearly any clinical data model. In this paper, we describe how Leaf can be incorporated into existing enterprise workflows and infrastructure, allowing clinicians and researchers a window into clinical data while simultaneously reducing the informatics burden of supporting similar tools. As Leaf is now an open-source software project, we welcome discussion and collaboration and encourage other members of the CTSA community to join us in using and improving the tool.

## Acknowledgments

We gratefully acknowledge: our clinical and research users at the University of Washington for their suggestions and patience in testing Leaf; Robert Meizlik and Xiyao Yang for their development work on early versions of Leaf; Matthew Bartek (University of Washington); Nora Disis, Carlos De La Peña, Jennifer Sprecher, Marissa Konstadt (Institute of Translational Health Sciences); Sarah Biber and Melissa Haendel (Oregon Health and Science University, Center for Data to Health); and Shawn Murphy, Isaac Kohane, Diane Keogh, Douglas MacFadden and other members of the i2b2 and SHRINE teams whose groundbreaking work we have been inspired by and remain indebted to.

## Funding

This work was funded by the National Institutes of Health, the National Center for Advancing Translational Sciences, as well as NIH/NCATS UW-CTSA grant number UL1 TR002319 and NIH/NCATS CD2H grant number U24 TR002306.

## Competing Interests

None.

